# Behavioral signatures of a developing neural code

**DOI:** 10.1101/856807

**Authors:** Lilach Avitan, Zac Pujic, Jan Mölter, Michael McCullough, Shuyu Zhu, Biao Sun, Ann-Elin Myhre, Geoffrey J Goodhill

**Author notes:** Edmond & Lily Safra Center for Brain Sciences, Hebrew University, Jerusalem 9190401, Israel (Lilach Avitan).

## Abstract

During early life neural codes must develop to appropriately transform sensory inputs into behavioral outputs. Here we demonstrate a direct link between the maturity of neural coding in the visual brain and developmental changes in visually-guided behavior. In zebrafish larvae we show that visually-driven hunting behavior improves from 4 to 15 days post-fertilization, becoming faster and more accurate. During the same period population activity in the optic tectum refines, leading to improved decoding and information transmission of spatial position, particularly in the representation of the frontal visual field. Remarkably, individual differences in decoding can predict each fish’s hunting success. Together these results show how the neural codes required to subserve a natural behavior emerge during development.

## Introduction

During early development the brain must build neural codes appropriate for survival (Avitan and Goodhill, 2018). Techniques such as fluorescent calcium imaging for recording the activity of many neurons simultaneously have recently facilitated new insights into the development of neural coding. In the mammalian cortex in particular knowledge is expanding rapidly regarding the development of neural circuits (Levelt and Hübener, 2012; Ko et al., 2013; Smith et al., 2015; Chevée and Brown, 2018). However the behavioral repertoire in the neonatal period of most of the mammalian species studied in this regard is relatively limited (Brust et al., 2015). Therefore the impact of these early developmental changes in neural coding on measurable aspects of behavior remains largely unknown.

In contrast, zebrafish larvae display sophisticated natural behaviors from only a few days post-fertilization (dpf). Retinal ganglion cell axons reach the optic tectum (the main visual processing centre in zebrafish, homologous to the superior colliculus in mammals) from 2 dpf (Kita et al., 2015), and from 5 dpf their visuomotor system is already mature enough for them to hunt fast-moving prey such as *Paramecia* (Borla et al., 2002; Portugues and Engert, 2009; Muto et al., 2013; Portugues et al., 2014). A hunting event proceeds via a series of stereotyped bouts of movement driven by sequentially refined localization of the prey in visual space (McElligott and O’Malley, 2005; Marques et al., 2018). As zebrafish approach maturity, hunting behavior refines from bouts to much smoother movement (Westphal and O’Malley, 2013), however how it changes over the first few days of hunting experience remains uncharacterized. During this period it is possible to image the activity of large populations of neurons in the zebrafish brain using fluorescent calcium indicators. Spontaneous activity in the tectum over this time follows a specific developmental trajectory, showing greatest exuberance at 5 dpf and then refining (Avitan et al., 2017; Pietri et al., 2017). Tectal activity can be used to decode the spatial position of prey-like spots (Avitan et al., 2016). However whether the quality of this decoding changes with age, and how these developing neural properties are related to changes in behavior, remain unknown.

Here we show that there is a systematic refinement in hunting behavior over the first few days when zebrafish larvae begin to hunt prey, and that this is correlated with a systematic improvement in the quality of decoding and information transmission of spatial position in the optic tectum. The latter improvement follows a specific spatiotemporal trajectory, suggestive of a changing balance of priority between predator avoidance and prey capture. Furthermore, the hunting success of individual fish can be predicted from the quality of their tectal codes.

## Results

### Hunting performance improves over development

To study the development of a natural behavior, we imaged larval zebrafish hunting *Paramecia* at 5, 8–9 and 13–15 dpf (n=34, 22,33 fish respectively, different fish for each age, for brevity we hereafter refer to the older groups as simply 9 dpf and 15 dpf respectively). Zebrafish transition from self-feeding via the yolk sac to feeding via hunting behavior at 5 dpf (Kimmel et al., 1995). A hunting event commences with eye convergence and larvae maintain their eyes in a highly converged position for the duration of the event until they strike at the prey or abort the event (Bianco et al., 2011; Muto et al., 2013; Trivedi and Bollmann, 2013; Bianco and Engert, 2015).

Each larva was placed in a dish with 30–50 *Paramecia* and behavior was imaged for 15–25 mins using a high speed camera. A typical hunting event is shown in Figure 1A. We analysed all hunting events per fish and examined their developmental trajectory (see Methods). Hunting event rate increased over development (Figure 1B), (unless otherwise stated, in all figures each point represents the mean value for one fish over all events). Hunting events in which the fish struck toward a *Paramecium* became shorter in duration (Figure 1C), consisted of fewer bouts (Figure 1D), and increased bout rate (Figure 1E) over development, suggesting that larvae become more efficient at executing these behaviors. Since individual fish can develop at different rates we also examined these measures as a function of length in addition to age (Figure S1A), and all trends were preserved (Figure S1B-E).

**Figure 1:**
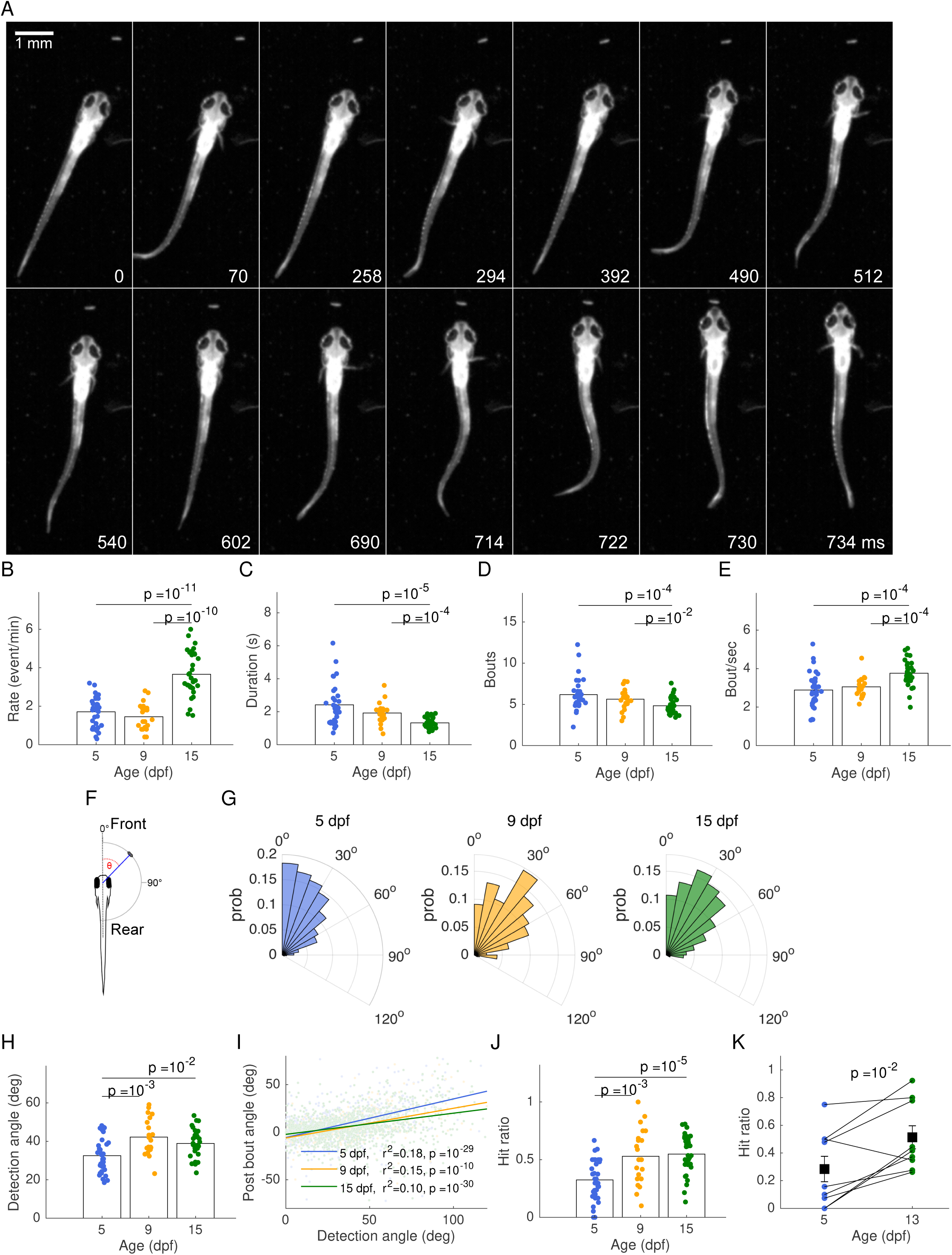
Hunting performance improves over development. A: A sequence of frames during a hunting event from the moment of prey detection (0 ms), showing orientation and eye convergence (70 ms), approach (294, 512 ms), strike (714 –730 ms) and prey capture (734 ms). B: Hunting event rate is higher for 15 dpf compared to 5 and 9 dpf fish. C: Hunting events become shorter over development. D: Number of bouts per event is lower for 15 dpf compared to 9 and 5 dpf fish. E: Bout rate is higher at 15 dpf compared to 5 and 9 dpf. F: Detection angle is defined as the angle between a *Paramecium* and the midline of the fish just prior to eye convergence. G: Distributions of prey detection angle for 5, 9 and 15 dpf fish (left, middle, and right respectively) change over development (5 vs. 9, *p* = 10^−7^; 5 vs. 15, *p* = 10^−6^; 9 vs. 15, *p* = 0.05; Two-sample Kolmogorov-Smirnov test). H: 15 and 9 dpf fish detect *Paramecia* slightly more toward the rear visual field compared to 5 dpf fish. I: The relation between prey detection angle and angle to prey after the first bout (here each point represents a bout rather than a fish). The slope decreases over development, indicating a more accurate alignment toward the prey. J: Hunting performance, as measured by the proportion of all hunting events per fish which are successful, increases over development. K: Hunting performance of individuals imaged at 5 and later at 13 dpf increases over development. Black squares represent the mean and bars SEMs

To examine the spatial relationship between the larva and the *Paramecium* of interest at the time of prey detection we calculated the angle between them at the frame prior to eye convergence (Figure 1F) (see Methods). These angles were predominantly in the frontal visual field (Figure 1G), with a slight increase in angle in older fish (Figure 1G, Figure S1F). The first bout tended to undershoot the angle required to align the fish with the prey, but this undershoot became smaller with age (Figure 1I).

By calculating a handedness measure (*R* − *L*)/(*R* + *L*), where *R* and *L* are the number of detections by the right and left eye respectively, we found no lateralization in *Paramecia* detection (Figure S1G,H). A hunting event was labelled as a hit event if the fish struck accurately at the *Paramecium* of interest (see Methods). The hit ratio i.e. the fraction of hit events out of all hunting events recorded per fish, increased over development (Figure 1J, Figure S1I).

For the results so far each fish was only evaluated at one age. For a smaller number of fish (n = 9) we performed the more challenging experiment of evaluating hunting performance in the same fish at both 5 and 13 dpf (see Methods). For 8 of 9 fish the hit ratio improved (Figure 1K), as did other measures of performance (Figure S2), demonstrating that improvement at the group level was reflected in improvement in individual fish. Thus, although hunting behavior is often considered to be highly stereotyped, its characteristics change substantially over early development, with more efficient execution of hunting events and improved hunting performance.

### Neural responses to frontal stimuli increase over development

We next asked whether these developmental changes in hunting behavior are correlated with developmental changes in the neural representation of the visual world. We embedded 4, 5, 8–9, 13–15 dpf fish (n = 11, 17, 10, 7 fish respectively, different fish for each age, and similarly to the above the older groups are referred as 9 dpf and 15 dpf respectively) in agarose for 2-photon calcium imaging of the optic tectum, a structure essential for hunting behavior (Gahtan et al., 2005) (see Methods). Of these, 17 fish had previously been imaged for behavior and are included in Figure 1. We recorded 61.6 min of evoked activity in response to twenty repetitions of nine 6° diameter spots (a size likely to be interpreted as prey (Bianco et al., 2011)) at azimuthal angles from 45° to 165° relative to the fish’s midline in its visual field (spots at smaller azimuthal angles failed to elicit robust responses). Spots were presented for 1 s each with a 19 s gap between spots, with a total of 180 spot presentations per fish (Figure 2A). Consistent with broad tectal receptive fields previously reported (Sajovic and Levinthal, 1982; Niell and Smith, 2005; Romano et al., 2015), each spot in the visual field elicited a response in a population of many cells. These population responses roughly matched the topographic retinotectal organization whereby spots in the frontal visual field elicited neural responses in the anterior tectum (Figure 2B,C).

**Figure 2:**
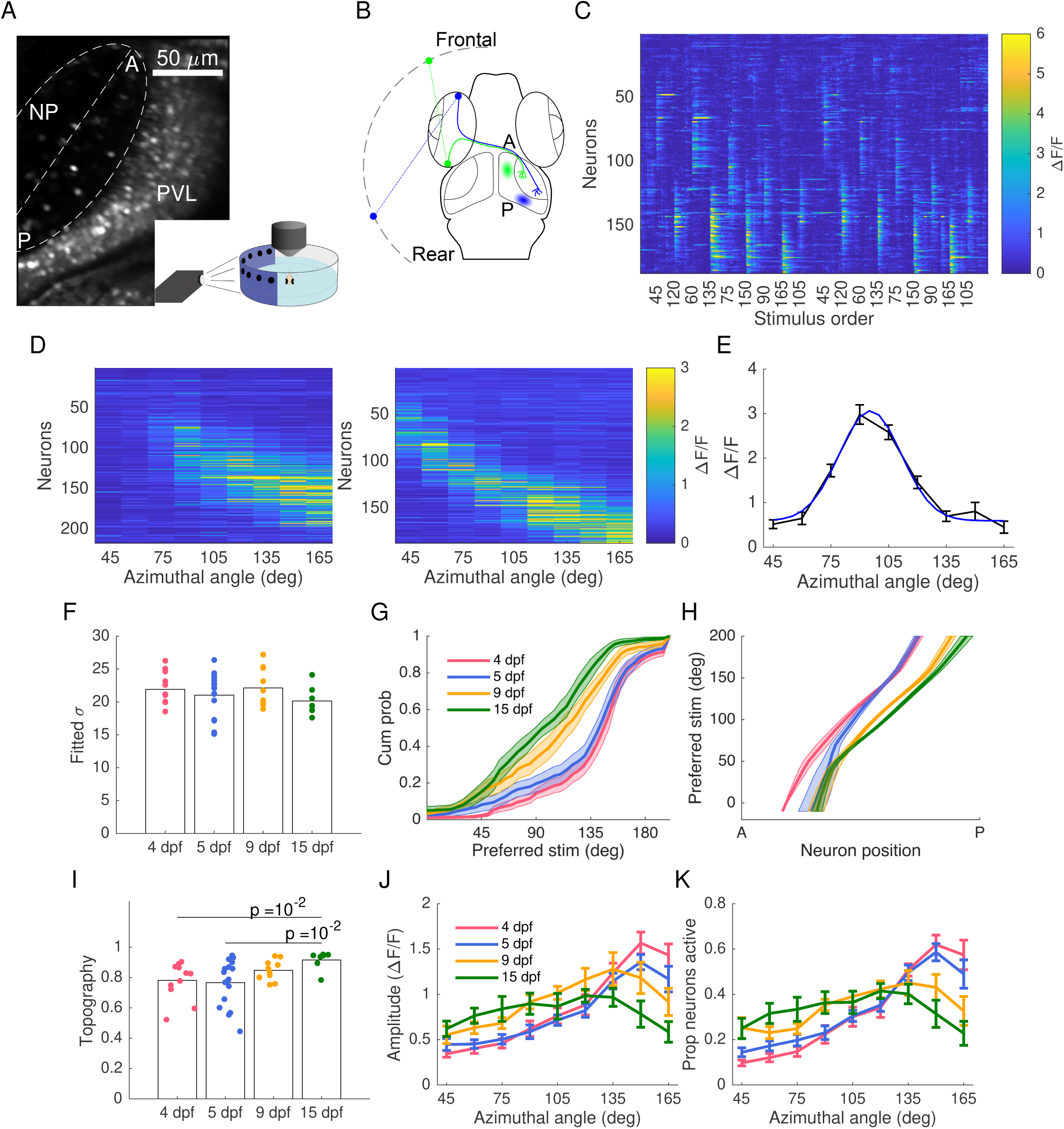
Changes in neural representation of the frontal visual field. A: An example tectal image from a 15 dpf fish. The neuropil (NP) contour of each fish was fitted with an ellipse with the major axis defining the tectal anterior–posterior (AP) axis. Periventricular layer (PVL), NP, anterior (A) and posterior (P) ends of the tectum are indicated. Inset: larvae were embedded in agarose with one eye facing the projected image for 2-photon calcium imaging. We recorded evoked activity in response to 20 trials of the stimulus set consisting of spots at positions 45°, 60°, 75°, 90°, 105°, 120°, 135°, 150°, 165° of the visual field where 0° was defined as the body axis. B: Schematic of retinotectal projection showing temporal retinal ganglion cells (green, representing the frontal visual field) projecting to anterior tectum, and nasal ganglion cells (blue, representing the rear visual field) projecting to posterior tectum. C: Population response of 187 PVL neurons from the fish shown in A elicited by two trials of visual stimuli. Neurons are sorted by their position on the AP axis. D: Raw receptive fields of 217 and 187 PVL neurons in an example 4 dpf (left) and 15 dpf (right) fish respectively. Neurons are sorted by their position on the AP axis and display a rough topography. E: Raw (black) and fitted (blue) receptive fields of an example neuron from the fish shown in A and C. F: Mean receptive field width did not change over development. G: More neurons were tuned to the rear visual field at 4 dpf, but this gradually balanced over development. Shading represents SEMs. Comparing the area underneath the curves for individual neurons grouped by age 4 vs.5 ns; 4 vs. 9 *p* = 0.01; 4 vs. 15 *p* = 10^−3^; one-way ANOVA with Bonferroni multiple-comparison correction. H: Neuronal selectivity shifted over development, so that at each tectal position neuronal tuning moved in a frontal direction. Shading represents SEMs. A linear fit to these curves showed similar slopes but different intersection points with the x-axis, indicating maps are shifted (4 vs. 15 *p* = 0.02, for slopes, and *p* = 0.04 for intercepts t-test). I: Topography improved over development (two-sample t-test). Variability also decreased over development (4 and 5 vs. 9 and 15, *p* = 0.008, two-sample F-test). J: At 4 dpf the population response amplitude was stronger in response to the rear visual field, but this balanced over development to give a more even response amplitude to stimuli covering the visual field. K: At 4 dpf a larger proportion of neurons were active in response to stimuli in the rear visual field, but again this balanced over development.

The receptive field of each neuron was calculated by averaging its response amplitude over stimuli repetitions (Figure 2D). We fitted each neuron’s raw receptive field with a Gaussian function with baseline offset (See Methods) (Figure 2E), and the angle at which the Gaussian peaked was defined as the neuron’s preferred stimulus. There was no change in receptive field width over development (Figure 2F). At 4 dpf the proportion of neurons tuned for the frontal visual field was much lower than for the rear visual field; however this tuning bias balanced over development with roughly even representation of the visual field by 15 dpf (Figure 2G). We then examined the relation between the position of the neurons on the anterior-posterior (AP) axis of the tectum and their tuning, and observed a shift of the tectal map with age (Figure 2H), this was not due to a change in eye position over development (Figure S3A-B). We further quantified topography as the correlation coefficient per fish between the preferred stimuli of neurons and their position on the AP axis. Topography was more precise at 15 dpf compared to 4 dpf and showed decreased variability for 9 and 15 dpf combined compared to 4 and 5 dpf combined (Figure 2I).

In addition to functional changes at the single-neuron level, there were also substantial changes in the statistics of the neural response at the population level. Response strength in the rear visual field was much higher compared to the frontal visual field at 4 dpf. However over development the strength of rear responses decreased while the strength of frontal responses increased (Figure 2J). There was a similar pattern for the proportion of neurons active (Figure 2K). Thus tectal representation of the visual field exhibits substantial spatial and functional changes over development, with a large increase in response strength to frontal stimuli.

### Decoding performance in the frontal but not rear visual field improves over development

The results so far demonstrate that tectal circuits undergo substantial functional changes at the single neuron and the population level between 4 to 15 dpf. We next asked whether these circuitry changes are reflected in the performance of decoding spatial information, and furthermore whether they are correlated with hunting performance. Using a linear decoder we quantified the relation between the original stimulus and the decoded stimuli position (Figure 3A-D) (results were qualitatively similar for other decoders (Figure S4A). Decoding performance in older fish was higher when averaged across the entire visual field compared to younger animals (Figure 3E). This improvement was due to improvement in the frontal visual field: mean decoding performance for frontal stimuli averaged over fish increased over development (Figure 3F). By 15 dpf performance was high and uniform over the entire visual field (Figure 3G). Notably, the improvement in decoding occurred despite there being no sharpening of receptive fields (Figure 2F).

**Figure 3:**
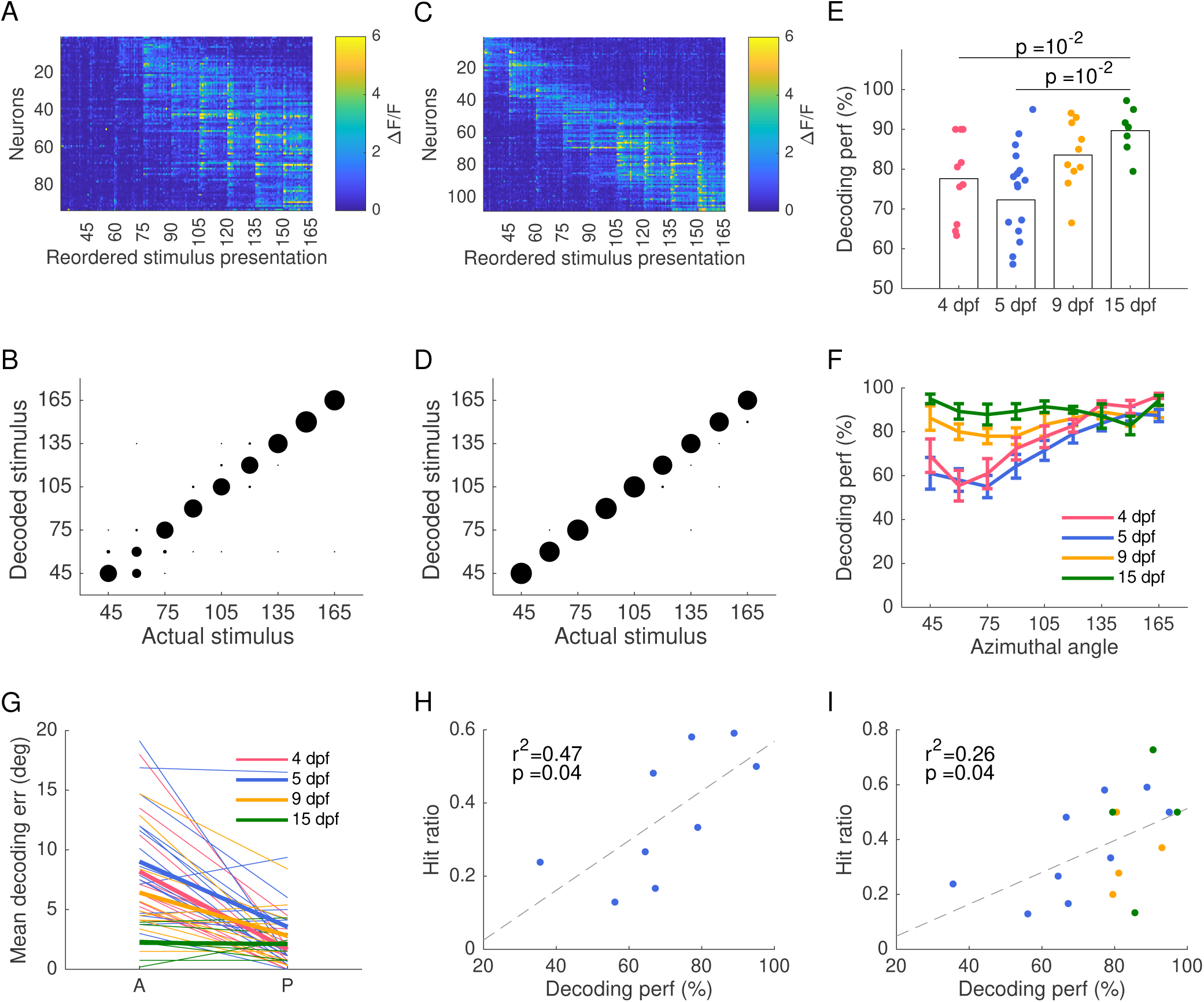
Decoding of stimuli in the frontal visual field improves over development. A: Population responses grouped and reordered by the stimulus presented in an example 4 dpf fish. B: Decoded stimuli for each of the 20 presentations of each stimulus as shown in A. Dots size is proportional to the fraction of presentations which decoded a particular stimulus value. C-D: Same as A-B for an example 15 dpf fish, showing better performance than the 4 dpf fish. E: Decoding performance over all stimuli per fish improved over development (t-test). F: Decoding performance became more even across the visual field with age. Error bars are SEMs. G: Decoding error for the most anterior two spots and the most posterior two points for each fish (thin lines), and the averaged error per age (thick coloured lines) at different ages showed better decoding performance of rear visual stimuli at 4,5, and 9 dpf (4 dpf, *p* = 10^−3^; 5 dpf, *p* = 10^−3^; 9 dpf, *p* = 10^−2^; paired t-test) but no difference at 15 dpf. H: Hit ratio for individual 5 dpf fish is correlated with their decoding performance, *p* = 0.07. I: Hunting hit ratio of individual fish including 5, 9 and 15 pdf is correlated with their decoding performance, *p* = 0.06.

There was substantial individual variation in decoding performance within an age group, particularly at 5 dpf when zebrafish are normally just beginning to hunt prey. But are these differences related to the ability of larvae to catch prey? We indeed found a correlation between decoding performance and hunting hit ratio per fish for the seven 5 dpf fish for which we had both behavioral and imaging data (Figure 3H). There was no dependence of hit ratio on fish length for the 5 dpf fish (Figure S4B), indicating that the correlation between hit ratio and decoding was not simply due to different levels of maturity within the 5 dpf population. However, this correlation was also present when fish from all ages for which we had both behavioral and imaging data were included (Figure 3I). Thus tectal decoding performance can directly predict the prey-capture abilities of individual fish, suggesting that individual differences in the ability to spatially localize prey constrain hunting performance.

### Information encoded in neural responses shifts and increases over development

To further investigate spatiotemporal changes in tectal coding, we next investigated how response entropy and mutual information were distributed across the tectum, and how these distributions changed with development. For this analysis we divided the stimuli into three subsets; the three frontal stimuli (45°, 60°, 75°), the three middle stimuli (90°, 105°, 120°) and the three rear stimuli (135°, 150°, 165°). We computed each neuron’s response entropy for each stimulus subset. Entropy showed localized increases on the AP axis consistent with the presence of a topographic organization (Figure 4A-C, left panels). We fitted the averaged entropy curves with a Gaussian function with baseline offset and examined peak entropy position. Consistent with an increasing proportion of neurons responsive to frontal visual stimuli (Figure 2K) there was an increase in the entropy for frontal stimuli over development (Figure 4A), and a shift towards a more posterior peak of entropy for all stimulus subsets in the tectum (Figure 4A-C, right panel), consistent with Figure 2H.

**Figure 4:**
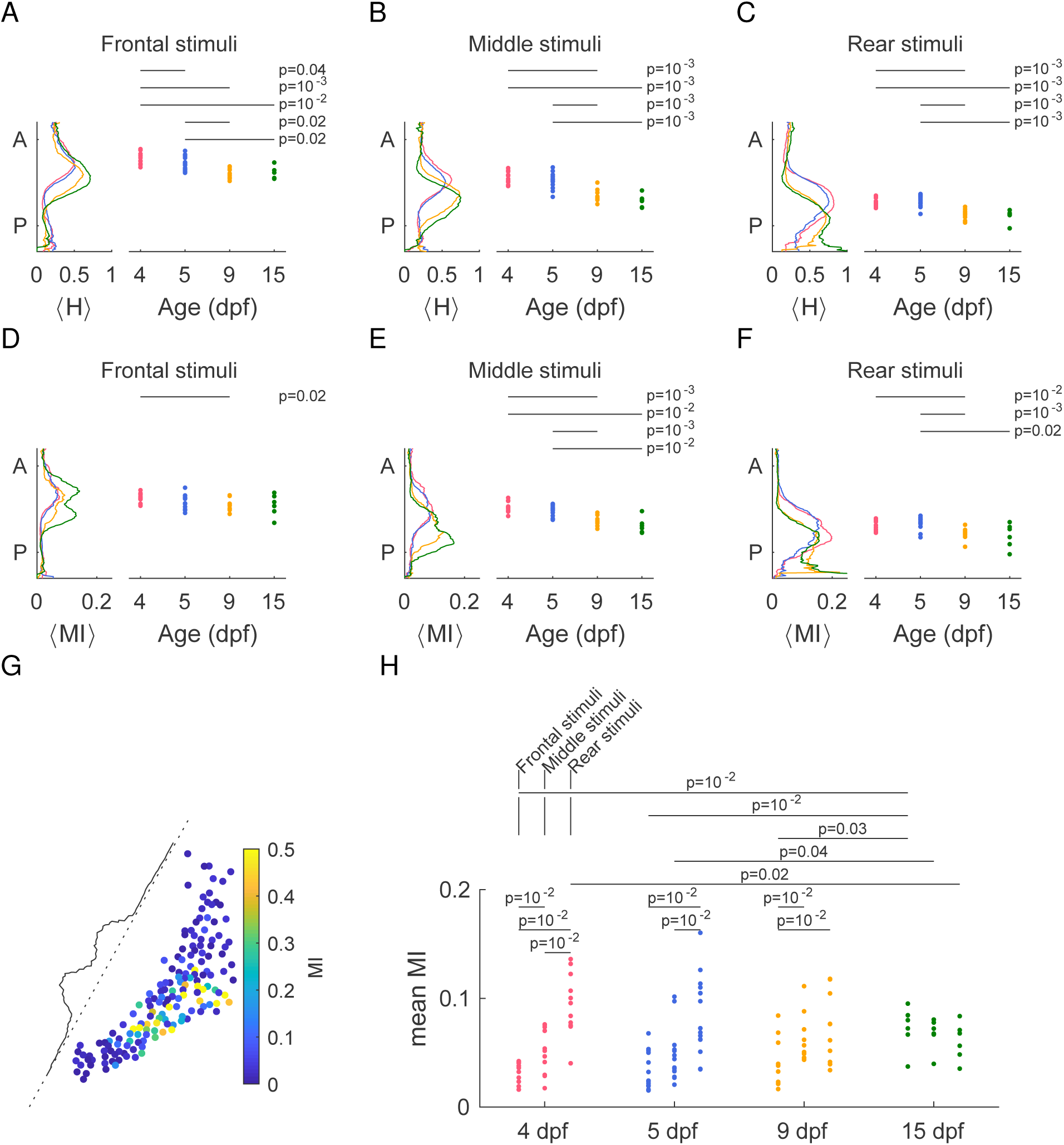
Information for the frontal visual field increases over development. A-C: Average entropy distribution along the AP axis for the three stimulus sets (left), and the respective peak position of the fitted curves (right) indicate a spatial shift in information over development for all stimulus sets. D-F: Mean mutual information distribution for the three stimulus sets along the AP axis (left) and the distribution of centre of mass (right) show a posterior shift in mutual information over development. G: Example of the distribution of mutual information for the three middle stimuli across the optic tectum of a 9 dpf fish, and the respective mean MI along the AP axis. H: Mean mutual information for the three stimulus sets over development. Mean mutual information increases over development for the three most frontal stimuli. All tests Wilcoxon rank-sum.

For each neuron we then computed the mutual information between the neuron’s response and the different stimulus subsets (example shown in Figure 4G), and examined the average mutual information along the AP axis (Figure 4D-F, left panels). The centre of mass of these distributions shifted posteriorly in the tectum for all stimulus subsets (Figure 4D-F, right panels) indicating a shift in stimulus information. However, this mutual information distribution along the AP axis was quite variable, and in some cases appeared bimodal (Figure 4D-F, right panels, example shown in Figure 4G, see also Figure S5A). To examine the source of this variability we considered a model of Bernoulli neurons with Gaussian receptive fields and tuning that tiled the visual field (Figure S5B, see Methods). The model allowed us to relate the mutual information between the activity of single neurons and the stimuli to the variance of the neurons’ response probability to the different stimuli, and suggested that high mutual information values are achieved for neurons with high variability (Figure S5C,D). We then used the model to examine the distribution of mutual information for a population of neurons, and showed that as receptive field size was increased, the local unimodal distributions of mutual information around each stimulus position merged to a global bimodal distribution (Figure S5E,F). This bimodal distribution was observed even when the map was irregular (Figure S5G, see Methods) consistent with experimental results (Figure S5A, middle). Thus, the bimodality seen in our data (Figure S5A) can be straightforwardly explained by a simple computational model.

Finally, we considered the mean mutual information between the different stimulus subsets over all neurons (Figure 4H). At 4 and 5 dpf information about rear stimuli was higher than frontal stimuli, consistent with higher decoding performance of rear stimuli compared to frontal stimuli at these ages. Over development there was an increase in mean mutual information for the frontal stimuli, consistent with improved decoding performance of frontal stimuli. Together, these results show that stimulus-related information increased over development overall, but that this increase was restricted to the frontal visual field.

## Discussion

By analysing behavior over a period in which zebrafish are learning to hunt fast-moving prey we found that their hunting performance improves, indicating that this is a highly dynamic period in their development (Figure 1). We then showed that these improvements can be linked to improvements in sensory coding (Figure 2-4). While increasing levels of fine motor control also likely contribute to the increase in hunting effectiveness (for instance changes in fin and body morphology can influence the generation of locomotor forces (McHenry and Lauder, 2006) and fish hydrodynamics (Müller and van Leeuwen, 2004), our results demonstrate a very clear link between hunting efficiency and coding efficiency.

We showed that over development an increasing proportion of tectal neurons become responsive to stimuli in the frontal visual field (Figure 2G), the tuning of tectal neurons at each point on the AP axis moves towards more frontal stimuli (Figure 2H), decoding of spatial position improves for frontal stimuli (Figure 3F), and the overall topography of the map improves (Figure 2I). The temporal region of the retina, which is stimulated by objects in the frontal visual field and innervates the anterior part of the tectum (Figure 2C), is the last retinal region to differentiate at 5-7 dpf (Patterson et al., 2013). This retinal temporal region exhibits a high density of cones, likely providing high-acuity vision in the corresponding region of the visual field (Schmitt and Dowling, 1999; Zimmermann et al., 2018). Thus part of the improvement we observed in both hunting performance and tectal decoding of frontal stimuli could have a retinal origin.

While the retina grows in concentric annuli (Reh and Constantine-Paton, 1983), the optic tectum grows by adding new neurons at its posterior end (Boulanger-Weill et al., 2017; Boulanger-Weill and Sumbre, 2019). These neurons then migrate anteriorly in the tectum and become tuned to more frontal stimuli (Boulanger-Weill et al., 2017), consistent with our results. Projections from the tectum to premotor areas have some topographic organization (Helmbrecht et al., 2018), suggesting that a simple topography-based code could provide the substrate for visuo-motor transformations. However due to the highly dynamic nature of the tectal map (for instance we found frontal shifts in tuning of around 50° at each tectal position between 4 and 15 dpf (Figure 2H)), such a topography-based code would presumably also require matched plasticity in downstream structures. Topography-based codes for spatial position also have lower performance than more statistically-motivated codes (Avitan et al., 2016). Together, these findings suggest that tectal coding strategies for spatial localisation may be more sophisticated than topographic mapping alone.

An interesting developmental progression in our behavioral data is that fish become increasingly better aligned with the prey in the first bout after the hunting event is initiated. While this is consistent with a gradual improvement in prey localisation in the frontal visual field over development, an alternative explanation is that it is due to improvements in the motor system. However in both cases it remains to be explained why the turn is usually an undershoot, rather than errors being randomly distributed between overshoots and undershoots. This undershoot was previously observed in 5-7 dpf zebrafish by (Patterson et al., 2013), who found it was not a predictive effect since prey were equally likely to be moving away from as towards the midline of the fish. One possibility is that, assuming the energy required for the turn increases with the turn angle, a second corrective turn after an undershoot requires less summed energy for both turns than a second corrective turn after an overshoot.

One question raised by our work regards the role of eye convergence for hunting. A clear rationale for this at the final stage of prey capture, when the fish strikes from close range towards a target positioned directly in front of the fish, is that it provides depth information via binocular vision (Bianco et al., 2011). However this does not explain why eye convergence first occurs at the initial stage of a hunting event, when the prey is too distant and misaligned with the fish for binocular cues to be relevant. A possible rationale for initial convergence raised by our work is that it moves the image of the prey towards the nasal retina and thus towards the posterior tectum, where spatial decoding of position is initially superior (Figure 3F).

Despite multiple attempts we were unable to reliably release a 2-photon-imaged fish from agarose such that it remained viable for 2-photon imaging at an older age. However, our behavioural results showed that the increase in hunting performance observed in individual fish tracked between 5dpf and 13 dpf (Figure 1K, Figure S2) is consistent with the increase in hunting performance observed at the group level over development. This suggests that all the developmental changes we observed at the group level are likely reflected in developmental changes at the individual level.

We showed that individual differences in behavioral performance could be predicted by the quality of decoding from neural activity in the tectum (Figure 3F). Relating fMRI measurements in humans to individual differences in psychological traits is a rapidly expanding area (Dubois and Adolphs, 2016). For instance (Schwarzkopf et al., 2010) showed that variations in the size of primary visual cortex could predict individual differences in visual perception, and it has been demonstrated that individual functional connectivity profiles can predict fluid intelligence (Finn et al., 2015) and creativity (Beaty et al., 2019). However, as far as we are aware ours is among the first demonstrations that neural decoding can be used to predict behavior at an individual level (Honegger et al., 2019).

Due to its unique advantages, the larval zebrafish has proven to be a very effective model for addressing fundamental questions about brain function at the systems level. However we have shown that both behavior and neural coding in the optic tectum undergo significant refinement between 4 and 15 dpf. The larval fish brain at this stage is likely undergoing highly dynamic processes of development across many different systems simultaneously. It will be instructive in future work to determine to what degree neural mechanisms found in larvae generalise to adults.

## Methods

### Zebrafish

All procedures were performed with approval from The University of Queensland Animal Ethics Committee. *Nacre* zebrafish (*Danio rerio*) embryos expressing *elavl3:H2B-GCaMP6s* were collected and raised according to established procedures (Westerfield, 1995) and kept under a 14/10 hr on/off light cycle. Larvae were fed live rotifers (*Brachionus plicatilis*) daily from 5 dpf.

### Hunting behavior assay

89 individual fish larvae (34,22,33 fish aged 5,8–9,13–15 dpf respectively) were placed (singly) into a clear bottom recording chamber (diameter: 20 mm; depth: 2.5 mm; Cover-Well Imaging Chambers. Catalogue number 635031, Grace Biolabs), containing E3 embryo medium with fluid depth 2.5 mm and 30–50 *Paramecia* (*Paramecium caudatum*). Chambers were placed on a heating plate heated to 28.5°C (Fryer), illuminated from the bottom with an LED ring (LDR2-100SW2-LA, Creating Customer Satisfaction (CCS) Inc., Kyoto, Japan) and imaged with a stereoscope (Zeiss StereoDiscovery V8, 0.5X objective lens) equipped with a CMOS camera (Grasshopper GS3-U3-23S6M-C, Point Grey), at 100 fps. To achieve a faster imaging rate for a small number of fish, chambers were illuminated from the bottom with an infrared LED ring (850 nm, LDR2-100IR2-850-LA powered by PD3-3024-3-PI, CCS), from the top with a white light ring (LDR2-100SW2-LA, CCS Inc., Kyoto, Japan) and imaged with a CMOS camera (Mikrotron 4CXP, Mikrotron) at 300 fps (*n* = 7) or 500 fps (*n* = 13). A small set of chambers (*n* = 12, 16 fish age 5 and 9 dpf respectively) were imaged without a heating plate, but no statistical differences in hunting behavior characteristics (duration, detection angle, number of bouts, hit ratio) were observed except for a decrease in hunting event rate in the non-heated plates (t-test). Overall a total of 2569 hunting events were recorded.

### Imaging hunting behavior of individuals over development

To track the development of hunting behaviour for individual fish, GCaMP6s larvae were individually housed from 1 dpf in dishes with age-matched non-GCaMP fish (to allow the target individual to be identified). We then used the assay described above to image hunting behaviour of the GCaMP6s larvae at both 5 and 13 dpf.

### Analysis of hunting behavior movies

Movies were visually inspected and measures were extracted using ImageJ (Grasshopper movies) or StreamPix (Mikrotron movies). Results for 12% of the movies (10/86) were cross validated between 2–4 observers to ensure measurement consistency and minimize inter-examiner variability. The start and end of each hunting event were determined by eye convergence (Bianco et al., 2011). For each fish and each event we extracted the following measures characterizing event properties. Event duration was defined as the time between eye convergence and divergence. Detection angle was defined as the angle between the larva’s head direction and the vector defined by the point between the eyes on the midline and the *Paramecium* (Figure 1F). The *Paramecium* of interest was defined as the nearest *Paramecium* toward which the fish performed the first turn movement. We further measured the post-bout angle between the larva and the *Paramecium* after the first bout, number of bouts between the start and end of the event, and hunting hit ratio. To evaluate hit ratio we labelled every event using three labels (hit, miss and abort). In an ‘abort’ event, the larva chases but never lunges toward the prey. In a ‘miss’ event, the larva chases, inaccurately lunges toward the prey, and therefore misses the *Paramecium*. In a ‘hit’ event the larva chases, accurately lunges and captures the *Paramecium* of interest. This label also includes events where the *Paramecium* is caught by the larva and immediately released, since the prey was accurately localized in these cases. Hit ratio was calculated as the ratio of hit events out of all hunting events (hit, miss and abort) for a given fish. For each measure we obtained an average over all events per fish, and changes in the statistics of these averages over development were examined.

To assist in the extraction of behavioral measures we used custom image processing software written in MATLAB to analyse 25 of the movies. Movies were first pre-processed to create a foreground mask for the fish and the *Paramecium*, allowing us to remove background information in each frame. A Gaussian background model of each pixel was fitted over the set of every tenth frame of the movie. A pixel was classified as foreground in a given frame if its intensity was lower than 3 standard deviations from the pixel’s mean intensity or higher than 3 standard deviations from its mean. To obtain an initial estimate of the position of the fish in each frame, we used connected components analysis on the foreground mask to extract the largest component, which was a mask of the fish. We then cropped a 128 × 128 pixels tracking window around the centre of mass of the fish mask to reduce computational load for the next stage of the processing pipeline.

To compute the position and heading angle of each fish we tracked a point on the centre of the swim bladder and the midpoint between the eyes using a custom correlation filter approach based on the Minimum Output Sum of Squared Error (MOSSE) filter Bolme et al. (2010). To make tracking more robust, we trained a set of MOSSE filters using different features from the image. These features included the pixel intensity and an extended set of 31 histogram of gradients (HOG) features described in Felzenszwalb et al. (2009). To train the MOSSE filters we: (i) selected a set of 10 training frames; (ii) centred and rotated the fish to 0 degress in each frame; (iii) cropped a 128 × 128 pixel window around the fish; (iv) extracted the features to produce a set of training patches per feature; and (v) trained the MOSSE filter for each feature on its respective set of training patches. To enable tracking of the fish at different orientations, we repeated the filter training process for 72 evenly spaced rotations of the training patches over 360 degrees to create a rotated filter stack for each feature. To track the desired point on the fish in a given frame, we correlated the cropped tracking window with the rotated filter stack for each feature. We then computed the mean correlation over the set of features and identified the rotation that results in the maximally correlated pixel. This pixel was taken to be the location of the desired tracking point. The tracked trajectories for the eye midpoint and swim bladder were both smoothed using a Kalman Smoother (Rauch–Tung–Striebel algorithm) (Rauch et al., 1965) with Newtonian mechanics and constant acceleration.

To detect bouts we used the kinematics of the midline of the tail. The tail midline was estimated by performing a morphological thinning operation on the fish mask using MATLAB’s *bwmorph* function and pruning the resulting skeleton by finding the geodesic path between the swim bladder and the tip of the tail in the corresponding connected components graph. The midline was then smoothed in both *x* and *y* directions using a moving mean. A 1-dimensional constant velocity Kalman smoother was used to estimate the length of the tail which varies with the pitch of the fish. Ten evenly spaced points were then extracted by linearly interpolating along 95% of the length of the midline starting at the head end. We discarded the most posterior 5% of the tail because the relative illumination of this segment of the tail was low, resulting in noise. We then computed the angle *θ*_*i*_ between tail segments between the *i*-th and *i* + 1-th tail point for *i* = 1, 2, …, 9. Subsequently we computed the mean of the angular velocity 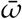 for the 4 most posterior tail segments:

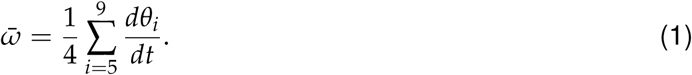

We then took the absolute value 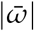 and applied a moving mean of 5 samples to smooth. Bouts were detected by applying a threshold which was selected manually and verified by visual inspection. We applied additional constraints that bouts must be at least 50ms long, and adjacent bouts must be separated by more than 20 ms else they were merged and treated as a single bout.

To track the location of *Paramecia* in each frame, the foreground mask was first filtered to retain connected components with area between 9 and 196 pixels. The centroids of each component were treated as detected locations of *Paramecia*. Each detected centroid was assigned to an existing track from the previous frame based on the minimum Euclidean distance between the detection and the existing track. If no suitable track existed then a new track was created. A constant velocity Kalman filter was used to track prey through short periods of occlusion such as when they swum under the fish.

Finally, each hunting event was manually inspected to record the beginning and end of the event, identify the *Paramecium* being hunted, and determine feeding event score. Using this information in conjunction with the fish and prey tracking enabled the computation of the behavioral measures. Results were manually validated and corrected as necessary using annotated output videos produced by the software.

### 2-photon calcium imaging

Zebrafish larvae were embedded in 2.5% low-melting point agarose, positioned at the centre of a 35 mm diameter plastic petri dish and overlaid with E3 embryo medium. Time-lapse 2-photon images were acquired at the Queensland Brain Institute’s Advanced Microscopy Facility using using a Zeiss LSM 710 inverted 2-photon microscope. A custom-made inverter tube composed of a pair of beam-steering mirrors and two identical 60 mm focal length lenses arranged in a 4f configuration was used to allow imaging with a 40X/1.0 NA water-dipping objective (Zeiss) in an upright configuration. Samples were excited via a Spectra-Physics Mai TaiDeepSee Ti:Sapphire laser (Spectra-Physics) at an excitation wavelength of 940 nm. Calcium imaging was performed at a depth of 70 µm from the dorsal surface of the tectal midline. Laser power at the sample plane ranged between 12 to 20 mW. Emitted light was bandpass filtered (500–550 nm) and detected with a nondescanned detector. Time-lapse images (416 × 300 pixels) were obtained at 2.2 Hz for 96.6 mins (the first 35 mins was spontaneous activity in the dark, which is not considered here). To improve stability of the recording, chambers were left to settle prior to imaging for 3 h.

### Visual stimulation

Visual stimuli were projected onto white diffusion paper placed around the wall of the petri dish, using a pico-projector (PK320 Optoma, USA), covering a horizonal field of view of 174°. To prevent stimulus reflection, the opposite side of the dish was covered with low-reflection black paper. To avoid interference of the projected image with the signal collected by the detector, a red long-pass filter (Zeiss LP590 filter) was placed in front of the projector. Larvae were aligned with one eye facing the projected side of the dish and body axis at right angles to the projector direction. Visual stimuli were generated using custom software based on MATLAB (MathWorks) and Psychophysics Toolbox (http://psychtoolbox.org). Each trial consisted of 6 degree diameter black spots at nine different positions from 45° to 165° with 15° intervals, with their order set to maximize spatial separation within a trial (45°,120°, 60°, 135°, 75°, 150°, 90°, 165°, 105°). 0° was defined as the direction of the larvae’s body axis. Spots were presented for 1 s each, followed by 19 s of blank screen. We projected 20 consecutive trials of nine spots with 5 s of inter-trial interval. A protocol composed of 20 trials of 11 spots (15°-165° with 15° interval) was used in one fish, and a protocol with 5 trials of 11 spots and 20 repetitions of 8 spots (60°-165° with 15° interval) was used in 12 fish. Edits to the experimental protocol were made since no anterior responses were detected early in development and we had no *a priori* knowledge this was a developmental effect.

### Tuning curves

For each neuron we calculated the mean amplitude across frames 4–7 post stimulus presentation. These amplitudes were then averaged per stimulus, providing for each neuron a curve of mean amplitude in response to all presented stimuli. Cubic spline interpolation was used to estimate the amplitude values between the presented stimuli at 5°intervals. This interpolated curve of amplitudes was fitted with a Gaussian with baseline offset. Fit starting points used the mean as the stimulus value eliciting response peak amplitude, and the initial value for standard deviation was varied from low to high values. The fitted curve which provided the highest goodness of fit (adjusted *r*^2^) was selected as the fit of the tuning curve. Neurons with goodness of fit greater than 0.7 were deemed to be selective neurons.

### Decoding stimulus identity

We used linear discriminant analysis to decode stimulus identity. The algorithm received as an input a set of population response vectors (20 × 9=180 vectors representing response averaged over frames between 1.5 s and 3.5 s post stimulus presentation) and their respective stimulus labels (i.e. the stimulus evoking each population response vector). The algorithm output the linear discriminant coefficients for the population for the nine different stimuli. Decoder classification coefficients were calculated from a training set separate from the test set to be decoded, using a leave-one-out strategy, in which a single population response vector was decoded using the statistics calculated on the basis of all presentations other than the test vector. Given a test population response, scores were calculated for each stimulus and the respective stimulus identity probability. The test population response was classified as the most probable stimulus. Overall decoding performance per fish was defined as the proportion of population responses classified correctly out of the 180 responses tested.

In addition to a linear decoder we used several other decoders. A topography-based decoder which relies on the Centre of Mass (CoM) of tectal responses as shown in (Avitan et al., 2016). By averaging the CoM of all the responses in the training set evoked by a given stimulus, we obtained a set of CoMs each corresponding to a particular stimulus. We then calculated the CoM of a given test response, and assigned it to the stimulus with the nearest CoM. A Maximum Likelihood decoder (ML) was used as shown in (Avitan et al., 2016), where decoding the population response consisted of searching for the stimulus, which had the highest probability of evoking a given population response. We then used three decoders taking into account only mean of the responses (M),or the mean and variance of the responses (M+V), and lastly the mean and the full covariance matrix (M+Cov). For the decoder based on mean responses, population responses in the train set were averaged per stimulus, resulting in a mean population response representing each stimulus. We then calculated the Euclidean distance between the test response and any of the mean responses, and assigned the test response to the stimulus whose corresponding mean response was closest. For the decoders based on either mean and variance or the mean and entire covariance matrix of the responses, we first calculated the covariance matrix. Similar to (Smith et al., 2015), the stimulus independent covariance matrix was computed as the mean of the covariance matrices for the individual stimuli

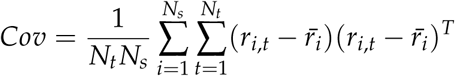

Where *N*_*t*_ is the number of trials, *N*_*s*_ is the number of stimuli, and *r*_*i,t*_ is the population response to stimulus *i* in trial *t*. The covariance matrix was used in two ways: as a whole for the M+Cov case, or where all of diagonal values were zeroed (M+V). Inspired by the discriminability measure introduced in (Smith et al., 2015), we evaluated the multivariate normal distributions centred at the mean responses to the different stimuli at the test response. These densities were normalized so that their value was 1 at the center. The stimulus providing the highest value at the test response was selected as the decoded stimulus.

### Information theory analysis

We binarized the raster of the neurons based on a threshold of mean amplitude plus two standard deviations for each neuron (Avitan et al., 2017). We then estimated the probability that a neuron was active in response to a particular stimulus *s, P*(*r* = 1|*s*), in the frames between 1.5 s and 3.5 s after any of the presentations of stimulus *s*. The probability of the stimulus, *P*(*s*), was uniform by experimental design. Mutual information between the activity of any particular neuron and the stimuli was then computed as where the first term is the entropy of that neuron. Besides the complete set of stimuli we sometimes considered the mutual information and entropy for different stimulus subsets. In these cases the stimulus sum was restricted to only those stimuli and *P*(*s*) was appropriately normalised.

The mutual information is trivially bounded by the smallest of either the stimulus entropy or response entropy. In our case, where the stimuli were approximately uniformly distributed on subsets of at least 3 stimuli, the stimulus entropy always exceeded the response entropy so that mutual information was bound by the latter. We obtained the average distribution of mutual information and entropy along the anterior-posterior axis using a moving average approach. For every point along the axis we computed the average distribution at this point as the mean across the neurons that were projected into a small interval of 0.15 times the length of the AP axis around this point.

### Modelling information theory results

We considered a stimulus variable *S* drawn from a discrete set of stimulus values where each value corresponds to a stimulus azimuthal angle 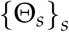. We modelled the activity *X*^*θ*^ of a single neuron tuned to angle *θ* as a Bernoulli variable with a response parameter modulated by the stimulus that is presented, i.e. 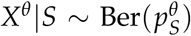, where 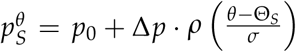 and *ρ* is an unnormalised Gaussian function centred at 0 with variance of 1, so that the width of the neuron’s receptive field is parameterised by *σ* and its preferred azimuthal angle is *θ*. (Figure S5B).

The mutual information between the stimulus variable and the activity of each neuron is

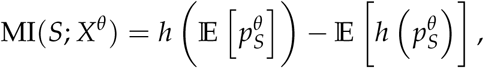

where *h*(*p*) = −*p* log_2_ *p* − (1 − *p*) log_2_ (1 − *p*) is the Bernoulli entropy.

Imposing beta distributions on the response probabilities, 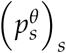, with shape parameters *α* and *β* we can explicitly compute the mutual information as

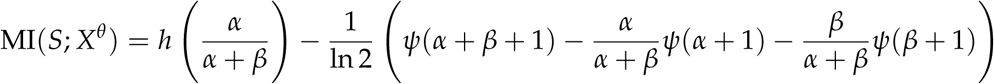

where *ψ* denotes the digamma function. Since the shape parameters can be uniquely expressed in terms of the expectation and mean of the corresponding distribution,

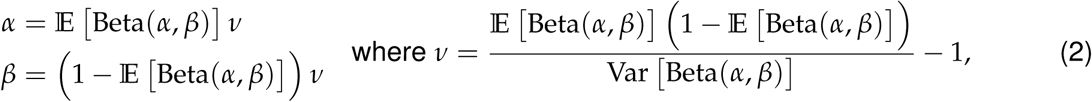

this yields an explicit expression of the mutual information in the Bernoulli model with beta distributed Bernoulli parameters. This is particularly instructive when considering the possible values of mutual information, parameterised in terms of expectation and variance of the response probabilities (Figure S5C,D). In order for a neuron to express large values of mutual information, the variance has to be large and consequently the expectation close to 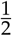 : for any random variable *Q* taking values in the unit interval, Var [*Q*] ≤ 𝔼 [*Q*] (1 − 𝔼 [*Q*]).

The dependence of mutual information on the variance can also be seen directly, if we express 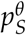 in terms of deviations from its mean and assume that these deviations are sufficiently small. The mutual information can then be expanded around 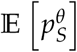 to yield

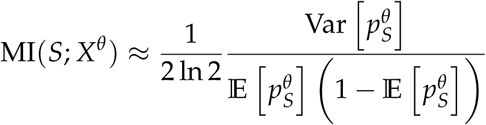

where errors to this approximation are cubic in the deviations from the mean.

We considered a population of neurons that are modelled by such a Bernoulli model and placed along a sensory axis representing the visual field, parameterised by angles from 0° to 180°. In the case of a regular map, the position of neurons along this axis corresponds to the centre of their receptive fields (their preferred azimuthal angle). To introduce irregularity into this sensory map we kept the position of the neurons constant but shifted the receptive field centres by adding Gaussian noise with mean 0 and standard deviation of 9°. We matched the parameters of the neuron model closely to the experimental data (*p*_0_ = 0.05, Δ*p* = 0.85) with three stimuli around the centre of the visual field (75°, 90°and 105°) and considered a population of 200 neurons. In order to describe a continuous mutual information distribution along the sensory axis we computed a moving average with sliding window width of 9°.

## Acknowledgements

We gratefully acknowledge funding from Australian Research Council grants DP170102263 and DP180100636. Imaging was performed at the Queensland Brain Institute’s Advanced Microscopy Facility using Zeiss LSM 710 2-photon microscope, generously supported by the Australian Government through the ARC LIEF grant LE130100078.

## Supplementary information

**Figure S1:**
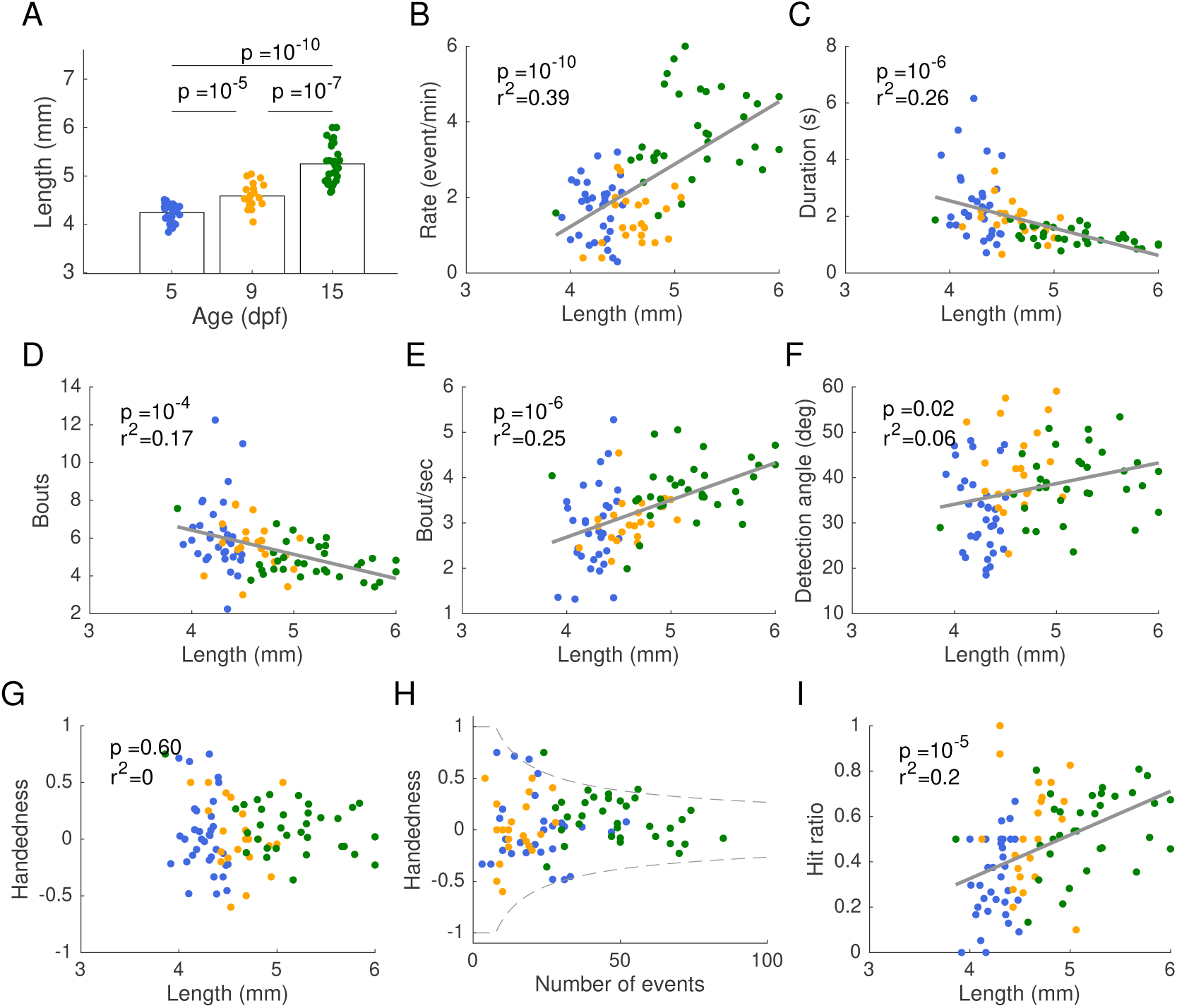
Changes in hunting behaviour as a function of fish length. A: Fish length increased over development, though with a substantial overlap between age groups. B: Event rate increased as a function of length (colours indicate age as shown in panel A). C: Event duration decreased as a function of length. D: Number of bouts decreased as a function of length. E: Bout rate increased as a function of length. F: Detection angle increased as a function of length. G: Handedness did not depend on length. H: Handedness decreased as a function of number of hunting events recorded. Handedness index mostly fell within the region expected under the null hypothesis that detection events are independent and uniform Bernoulli instances with a probability of at least 99% (marked by the grey lines), indicating no lateralization in *Paramecia* detection. I: Hit ratio increased with length.

**Figure S2:**
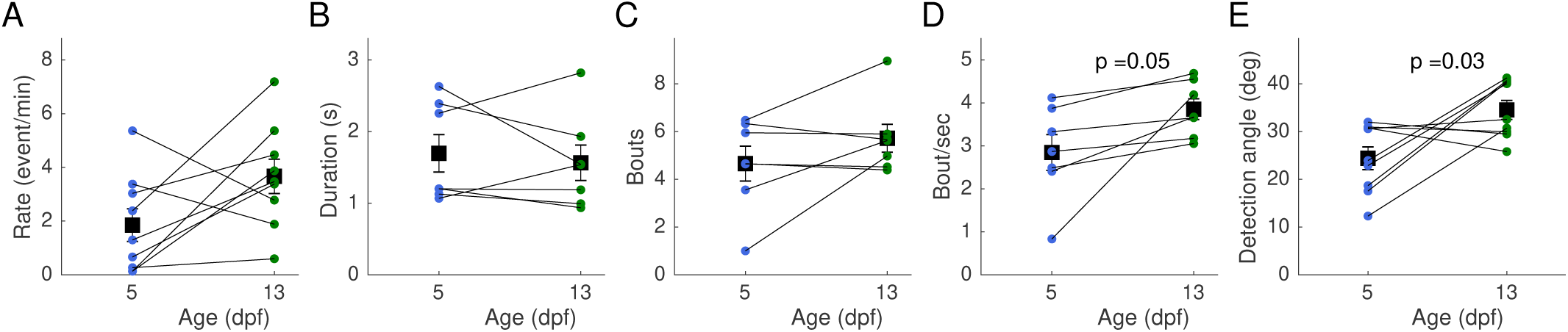
Hunting behaviour of individuals improve over development. A: Event rate shows an increase trend, similar to the effect shown when pooling fish. B: Hunting duration is not different, probably due to the low number of samples. C: Number of bouts per event shows an increase trend, similar to the effect shown when pooling fish D: Bout rate increases over development, similar to the effect shown when pooling fish. E: *Paramecia* detection angle increases over development, similar to the effect observed when pooling fish.

**Figure S3:**
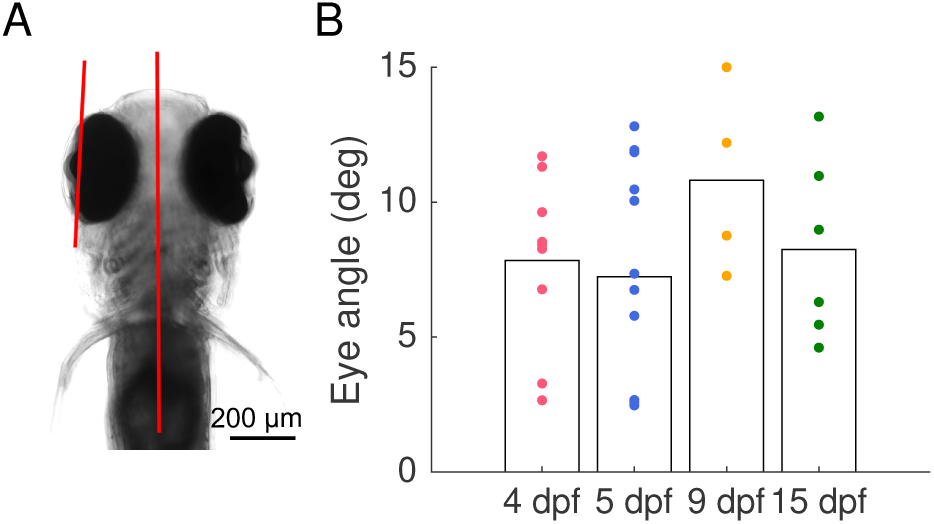
No difference in eye position during 2-photon imaging over development. A: The head of the fish was imaged at the end of every 2-photon imaging session and the angle between the eye and the midline was calculated. A: No difference in the angle between the eye and the midline during 2-photon experiments over development was observed.

**Figure S4:**
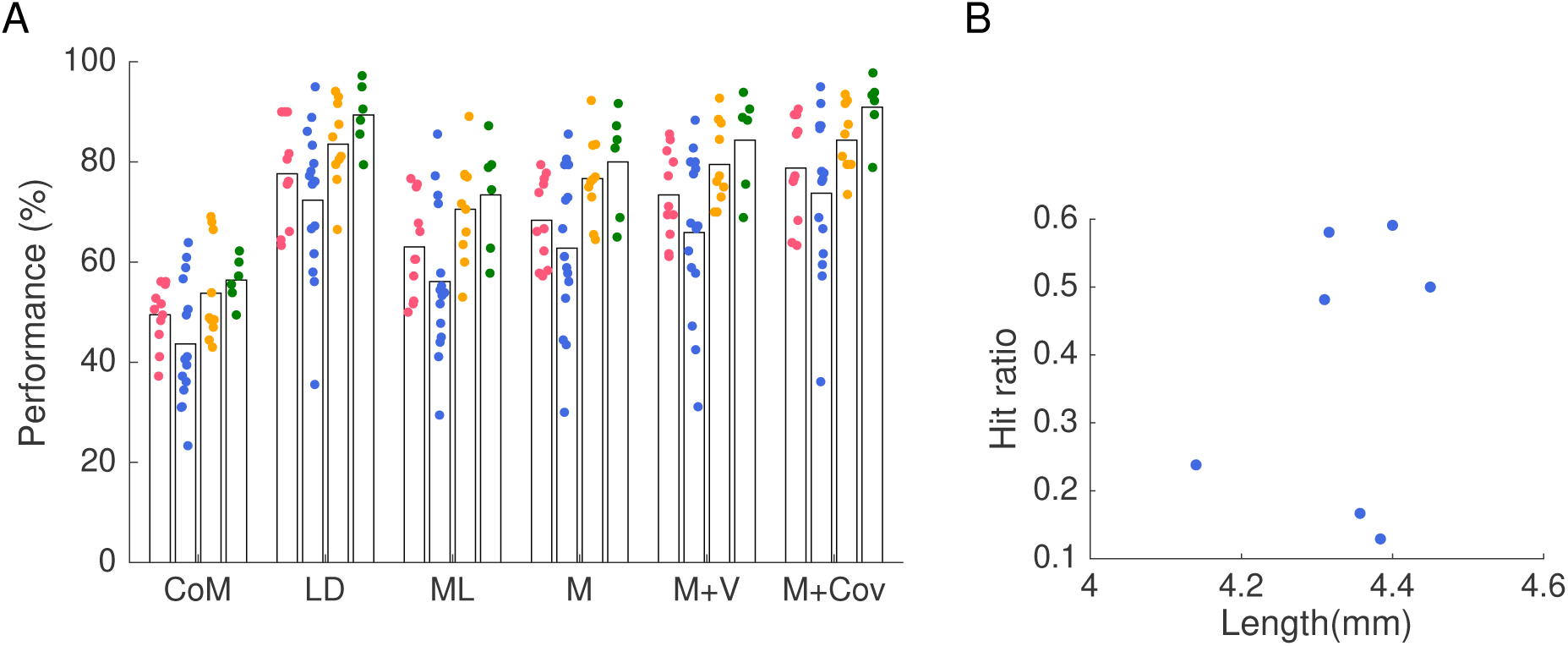
Comparison of different decoders. A: Decoding performance improves over development. Decoders used are: a topography based decoder which uses the response centre of mass (CoM), a Linear decoder (LD), a Maximum likelihood decoder (ML), a decoder which is based on population mean response (M), a decoder based on response mean and variance (M+V), and a decoder based on mean, variance and covariance (M+Cov). B: No (linear) relation between fish length and hunting hit ratio, *r*^2^ = 0.07, *p* = 0.6.

**Figure S5:**
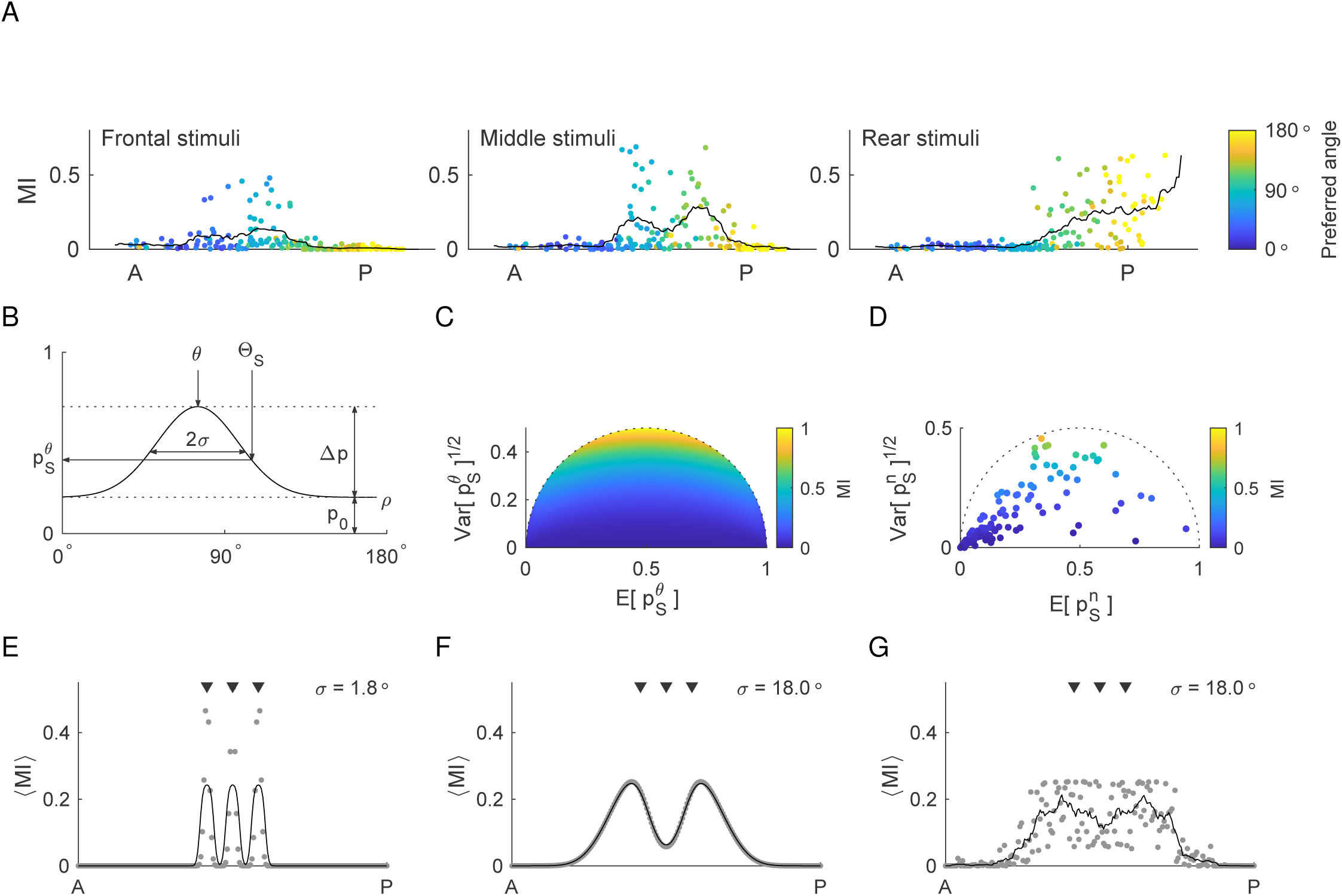
The distribution of mutual information over the AP axis appears bimodal for wide receptive fields. A: Distribution of mutual information between tectal neural response and the frontal, middle and rear stimulus sets along the AP axis in an example fish. Variations of bimodal distributions were apparent in all fish. Mutual information between neural activity and the middle stimulus set appears bimodal. Neurons’ preferred azimuthal angle is color coded. B: The response probability 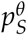 for a neuron in the Bernoulli model with a constant offset *p*_0_ and a term to modulate its response 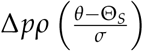. The neuron has a Gaussian receptive field function *ρ* of width *σ* and a preferred azimuthal angle *θ* and the stimulus angle is Θ_*S*_. C: Mutual information as a function of expectation and variance of beta-distributed response probabilities. Large values of mutual information can only be achieved for response probabilities that have a large variance. D: Mutual information as a function of expectation and variance of response probabilities estimated from tectal neural activity for the set of middle stimuli. Only very few neurons show high enough variance in their response probabilities to reach large values of mutual information. E–G: Simulated mutual information distribution, for a population of 200 neurons. The distribution is computed as a moving average of mutual information of the different neighbouring neurons. The triangles indicate the position of the stimuli at 75°, 90° and 105°. For narrow receptive fields and a regular sensory map, the distribution appears locally unimodal around the stimulus positions (E). For wider receptive fields which match experimental data and in the case of a regular sensory map (F) as well as an irregular sensory map (G), the mutual information appear globally bimodal due to overlapping receptive fields to the different stimuli.

